# A Generalizable Speech Emotion Recognition Model Reveals Depression and Remission

**DOI:** 10.1101/2021.09.01.458536

**Authors:** Lasse Hansen, Yan-Ping Zhang, Detlef Wolf, Konstantinos Sechidis, Nicolai Ladegaard, Riccardo Fusaroli

## Abstract

**Objective:** Affective disorders are associated with atypical voice patterns; however, automated voice analyses suffer from small sample sizes and untested generalizability on external data. We investigated a generalizable approach to aid clinical evaluation of depression and remission from voice using transfer learning: we train machine learning models on easily accessible non-clinical datasets and test them on novel clinical data in a different language.

**Methods:** A Mixture-of-Experts machine learning model was trained to infer happy/sad emotional state using three publicly available emotional speech corpora in German and US English. We examined the model’s predictive ability to classify the presence of depression on Danish speaking healthy controls (N = 42), patients with first-episode major depressive disorder (MDD) (N = 40), and the subset of the same patients who entered remission (N = 25) based on recorded clinical interviews. The model was evaluated on raw, de-noised, and speaker-diarized data.

**Results:** The model showed separation between healthy controls and depressed patients at the first visit, obtaining an AUC of 0.71. Further, speech from patients in remission was indistinguishable from that of the control group. Model predictions were stable throughout the interview, suggesting that 20-30 seconds of speech might be enough to accurately screen a patient. Background noise (but not speaker diarization) heavily impacted predictions.

**Conclusion:** A generalizable speech emotion recognition model can effectively reveal changes in speaker depressive states before and after remission in patients with MDD. Data collection settings and data cleaning are crucial when considering automated voice analysis for clinical purposes.

**Significant outcomes:**

- Using a speech emotion recognition model trained on other languages, we predicted the presence of MDD with an AUC of 0.71.
- The speech emotion recognition model could accurately detect changes in voice after patients achieved remission from MDD.
- Preprocessing steps, particularly background noise removal, greatly influenced classification performance.

**Limitations:**

- No data from non-remitters, meaning that changes to voice for that group could not be assessed.
- It is unclear how well the model would generalize beyond Germanic languages.

**Data availability statement:** Due to the nature of the data (autobiographical interviews in a clinical population), the recordings of the participants cannot be shared publicly. The aggregated model predictions and code used to run the analyses is available at https://github.com/HLasse/SERDepressionDetection.

## Introduction

Major Depressive Disorder (MDD) is a mental disorder affecting more than 163 million people worldwide ^1^, and comprises symptoms related to abnormalities in mood, cognitive ability, and psychomotor function ^2^. Current approaches to screening and monitoring of symptoms of depression primarily depend on self-reports, and are therefore often confounded by issues such as underestimating symptom severity, recency effects, and acquiescence ^3-5^. More efficient and objective screening and monitoring of depressive clinical features has the potential for relieving the disease burden significantly by scaffolding more timely and personalized interventions or by providing a measure of efficacy of the current treatment ^6^.

Using voice as a marker for depression is appealing as analysis and assessment can be done cheaply, remotely, and non-invasively. Depressed speech is characterized by a slow speaking rate, reduced inflection and prosody, and low volume ^7,8^. Correspondingly, a range of acoustic features have been identified as predictive of depression, ranging from changes to fundamental frequency (perceived as pitch) to more abstract spectral representations of speech ^9^. Multiple studies have used these acoustic features in machine learning models to discriminate depressed patients and healthy controls. Classification performance is highly varying, and accuracies between chance level and up to 94% are found in the literature depending on model and feature choice, dataset size, language, and validation method ^9,10^. However promising, it is still unclear how well these models would actually perform across a broader variety of clinical contexts.

For instance, a recent systematic review found that most of the studies predicting depression from voice did not fully evaluate the generalizability of these results to new data, i.e., they measured performance on the training sample or in a cross-validated fashion but not on held-out or external validation sets from different data sources or even just held-out test sets ^11-14^. This is probably motivated by small datasets being endemic to the field: high-quality clinical data is time-consuming and expensive to collect, and problematic to share. For instance, recent systematic reviews found that in related neuropsychiatric fields the median number of patients involved in studies of vocal markers were below 20 ^15-17^. Therefore, reducing the size of already small training samples by creating held-out data might be seen as problematic. However, even cross-validated performance has been shown to be unreliable on small or improperly nested datasets ^11,18(pp241-257),19^. It thus remains unclear how well these models would perform on new data collected in similar conditions.

Further, deploying automated voice analysis in clinical settings involves analyzing recordings of very heterogeneous patient populations collected in very heterogeneous physical settings with a wide variety of equipment. In other words, it involves generalizing the trained models to data that are potentially quite different from the original training set. We already know acoustic feature extraction from audio to be strongly influenced by background noise and recording conditions. For instance, one study has found consistent differences in the acoustic environments between their diagnostic groups, which could predict group membership with nearly the same performance as models based on acoustic features ^20^. Further, recent studies highlighted that acoustic patterns assumed to be characteristic of a given disorder might vary according to the language spoken^21^ and even the task employed ^16^. Therefore, it might not be enough to validate an algorithm even on held-out data collected in similar conditions to the training data, as this might produce overly optimistic evaluations of performance. It is clear that before clinical implementation is possible, generalizable models that perform well on data from other sources are needed and validation practices have to include more heterogeneity.

To overcome these limitations of small training data and lack of testing for external validity, we train a speech emotion recognition (SER) model and directly apply it to depression detection. Using a SER model for predicting depression is motivated by the findings that prosodic expressions of emotions are inhibited in depression and that experimental studies have found positive effects of adding SER to depression detection models ^22,23^. Datasets for SER are vastly more abundant, varied, and of higher quality than for depression assessment which is likely to produce more robust models. Further, by training a model solely for SER we are able to set aside our entire dataset of depressed patients and healthy controls for external validation thus providing a realistic view of generalizability.

## Aims of the study

The aims of the study are three-fold:

- *Aim 1:* to investigate the feasibility of using a generalizable emotion recognition model to directly predict depression,
- *Aim 2:* to assess the stability of these predictions over time as well as changes following remission from Major Depressive Disorder, and
- *Aim 3:* to quantify the effect of preprocessing steps such as background noise removal, speaker diarization (removal of speech from the interviewer), and time-window size on the quality and consistency of model predictions.

## Materials

### Speech Emotion Recognition Corpora and Model

The SER model was trained following Sechidis et al. ^24^ on the public CREMA-D ^25^, RAVDESS ^26^, and EMO-DB ^27^ datasets, all of which consist of recordings of sentences spoken by professional actors portraying different emotions. CREMA-D and RAVDESS include American English speech and EMO-DB German speech. A gradient boosted decision tree model was trained on each dataset separately to predict the probability of sounding happy or sad using Catboost ^28^ and combined in a Mixture of Experts (MoE) architecture ^29^ for ensemble prediction. MoE is a way of combining the predictions of multiple models (*ensembling*), which weights model predictions based on the distance of the input data to the constituent models’ training data. This way of ensembling allows one to train specialized models for e.g. different languages or recording conditions, and combine their knowledge at inference time hereby improving generalizability ^30^. In a previous study, using different SER corpora for external validation, the MoE model outperformed all constituent models as well as a single model trained on the pooled data from all the different corpora ^24^. Further details on feature extraction, training, and validation of the SER model are provided in the Appendix, hence the following sections in *Materials* and *Methods* relate to the depression corpus.

### Depressed Speech Corpus

The dataset used for depression assessment was collected at Aarhus University Hospital to investigate changes in social cognition in first-episode depression, and consists of 42 patients with first-episode MDD (two patients discarded due to missing data) and 42 healthy controls pairwise matched on gender, age, and educational level ^31,32^. Patients were recruited from general practitioners in the Central Denmark Region, and healthy controls were recruited via newspaper advertisements and offered monetary compensation if included in the study (1000 DKK). All participants were native speakers of Danish and met the following inclusion criteria: 1) first-episode major depression was the primary diagnosis, 2) the severity of depression was moderate to severe as measured by the 17-item Hamilton Rating Scale for Depression (HamD-17 > 17) ^33^, 3) patients were psychotropic drug-naïve, 4) no psychiatric comorbidity other than anxiety disorders. Physiological effects from psychotropic drugs can lead to changes in the voice ^34^, hence the inclusion of only psychotropic drug-naïve patients is crucial for the present study. Patients with head trauma, neurological illness, or substance use disorders were not permitted to the study. Exclusion criteria for healthy controls were the same as for depressed patients.

The dataset consists of audio recordings of the Indiana Psychiatric Illness Interview ^35^ conducted by a trained psychologist separately with each of the participants in Danish. Participants were asked to tell their life story in the first part of the interview, and in the second part to either reflect on their mental illness or on an emotionally distressing experience they have had in the last 2 years, depending on whether they were in the depression or control group. The interviews lasted between 20 and 50 minutes.

After the interview, the depressed patients underwent pharmacotherapy and individual psychotherapy. Those who entered remission within 6 months, defined as a HamD-17 score ≤ 7, were re-assessed with the same interview, along with the control group. As such, our dataset contains recordings of interviews with healthy controls at two timepoints six months apart (N=42/25), as well as interviews with 40 depressed individuals and a follow-up interview after six months with the subset who entered remission (N=25). Unfortunately, due to the focus of the original study, patients not in remission were not invited to the follow-up interview.

## Methods

**Table 1:** Demographic and clinical characteristics

|  | Visit 1 |  | Visit 2 |  |
| --- | --- | --- | --- | --- |
|  | Controls<br>(N=42) | Depression<br>(N=40) | Controls<br>(N=25) | Depression<br>(N=25) |
| **Demographics** |  |  |  |  |
| **Age** |  |  |  |  |
| Mean (SD) | 33.1 (12.3) | 32.5 (12.2) | 37.3 (12.5) | 30.9 (11.1) |
| Median [Min, Max] | 30.3 [18.1, 62.1] | 29.7 [18.8, 62.6] | 34.7 [19.7, 62.1] | 29.3 [18.8, 62.6] |
| Missing | 1 (2.4%) | 1 (2.5%) | 0 (0%) | 0 (0%) |
| **Gender** |  |  |  |  |
| Female | 33 (78.6%) | 31 (77.5%) | 21 (84.0%) | 20 (80.0%) |
| Male | 9 (21.4%) | 9 (22.5%) | 4 (16.0%) | 5 (20.0%) |
| **Education (years)** |  |  |  |  |
| Mean (SD) | 12.9 (1.94) | 12.3 (2.15) | 13.3 (1.97) | 12.7 (2.34) |
| Median [Min, Max] | 12.0 [10.0, 17.0] | 11.0 [9.00, 17.0] | 13.0 [11.0, 17.0] | 12.5 [10.0, 17.0] |
| Missing | 1 (2.4%) | 1 (2.5%) | 0 (0%) | 0 (0%) |
| **Clinical** |  |  |  |  |
| **HamD-17** |  |  |  |  |
| Mean (SD) | 1.61 (1.30) | 22.0 (3.47) | 1.78 (1.83) | 4.00 (3.10) |
| Median [Min, Max] | 1.00 [0, 5.00] | 22.0 [16.0, 30.0] | 1.00 [0, 5.00] | 3.00 [0, 11.0] |
| Missing | 1 (2.4%) | 1 (2.5%) | 2 (8.0%) | 4 (16.0%) |
| **Working Memory** |  |  |  |  |
| Mean (SD) | 6.78 (1.37) | 6.41 (1.37) | 6.68 (1.49) | 6.72 (1.28) |
| Median [Min, Max] | 7.00 [5.00, 9.00] | 7.00 [3.00, 9.00] | 6.00 [5.00, 9.00] | 7.00 [4.00, 9.00] |
| Missing | 1 (2.4%) | 1 (2.5%) | 0 (0%) | 0 (0%) |
| **Verbal Memory** |  |  |  |  |
| Mean (SD) | 9.61 (2.02) | 8.10 (1.64) | 9.72 (2.07) | 8.32 (1.35) |
| Median [Min, Max] | 10.0 [4.00, 12.0] | 8.00 [4.00, 11.0] | 10.0 [4.00, 12.0] | 8.00 [6.00, 11.0] |
| Missing | 1 (2.4%) | 1 (2.5%) | 0 (0%) | 0 (0%) |
| **Sustained Attention** |  |  |  |  |
| Mean (SD) | 0.924 (0.0527) | 0.888 (0.0658) | 0.929 (0.0538) | 0.885 (0.0721) |
| Median [Min, Max] | 0.934 [0.759, 1.00] | 0.888 [0.640, 1.00] | 0.943 [0.759, 1.00] | 0.886 [0.640, 1.00] |
| Missing | 1 (2.4%) | 1 (2.5%) | 0 (0%) | 0 (0%) |

**Note: HamD-17 = Hamilton Depression Rating Scale (17-item version), Working Memory (OTS) = One Touch Stockings of Cambridge test, Verbal Memory (HVL T-R) = Hopkins Verbal Learning Test-Revised, Sustained Attention (RVP) = Rapid Visual Information Processing test.**

### Data preprocessing

Background room noise, reverberation, and hum were removed from the audio recordings using iZotope RX 6 Elements ^36^. Long-term average spectra for each recording were inspected for possible noise artefacts and further cleaned if any were found.

To ensure that only voice segments from patients and controls were included, all interviews were manually segmented to only contain audio from the interviewee. We further removed all segments of audio without voice activity defined by a loudness threshold of −40dB and a minimum duration of 100 ms.

### Feature Extraction

From each audio recording, we extracted 13 mel-frequency cepstral coefficients (MFCC 0-12) with a window length of 25 ms and step size of 10 ms using the python_speech_features library ^37^. Mel-frequency cepstral coefficients (MFCCs) have been widely used in both speaker recognition ^38^, SER ^39^, and depression detection ^40^, and have several desirable properties such as being independent of the energy of the acoustic signal and robustness across genders ^41,42^. MFCCs represent movements of the vocal tract and are designed to mimic how the human ear perceives sounds by having high resolution in the lower frequencies and less in higher frequencies ^43^.

The zeroth MFCC was discarded as it represents the average energy of the signal. Though energy is often found to be a reliable vocal marker of depression, it is easily confounded by inconsistent recording conditions and its inclusion might therefore reduce the generalizability of the model.

The MFCC features were summarized in windows of different sizes (2, 5, 10, 15, 20, 25, and 30 seconds of speech as well as the whole recording) with 11 descriptive statistics: mean, variance, kurtosis, skewness, mode, IQR, percentiles 10th, 25th, 50th, 75th, and 90th. Summarization often results in better predictive performance than using raw features ^10,44^, and is far less computationally expensive. However, potential patterns that may emerge dynamically across segments of speech are disregarded. Windows of 30 seconds of speech were used for the main analyses.

Though other acoustic features such as pitch and energy are often found to be predictive of depression ^9^, we chose to focus on MFCCs as they have previously shown good predictive performance, can be robustly extracted, and are independent of energy and gender.

### Model Development

In order to assess whether SER predictions would discriminate depressed patients from controls (*aim 1*) and whether these predictions would be stable over time (*aim 2*), we built a Bayesian multilevel mixture model of the predictions. The SER model was applied to each speech sample from the depression corpus to obtain the log odds of sounding *happy* (1) or *sad* (0) for each time window, for each participant, at each visit. Given the heterogeneity of the symptom manifestations in depression ^45^, and only some patients entering remission, we adopted a mixture model of two gaussians to model the log-odds of sounding happy. In the mixture model, two Gaussian distributions best describing the data are estimated, and the probability of each speech sample to belong to one or the other can vary by participant - nested by diagnostic group as recommended in Valton et al. ^46^ to avoid pooling across groups - and visit. This also accounted for the presence of repeated measures, i.e., multiple predictions/time windows per participant. An interaction effect was used to assess the expectation that the depressed group should sound happier at the second visit as only those in remission were re-interviewed, while the voice of the control group should remain stable over time^1^.

To estimate the performance of the SER model for discriminating between depressed patients and healthy controls, the area under the receiver operating characteristic curve (AUC) using the raw predictions was calculated. Further performance metrics were calculated using the decision threshold that optimized AUC.

To more directly investigate *aim 2*, we also trained a hierarchical Bayesian classification model (logistic regression) to predict prognosis (remission yes/no) based on the depressed patients’ probability of sounding happy at the first visit. Baseline probabilities of sounding happy were modeled as varying by participant to account for repeated measures.

To assess the effects of the data preprocessing steps (*aim 3*), different datasets were created relying on the data from visit 1: a) data with background noise removal and without interviewer speech, b) data with background noise removal and including interviewer speech, c) data without background noise removal and including interviewer speech. How these preprocessing choices affected the difference in probability of sounding happy in patients and controls was assessed following the BEST approach suggested by Kruschke ^47^. Further, the area under the receiver operating-characteristic curve for classifying depression from controls was calculated for different window sizes (2, 5, 10, 15, 20, 25, and 30 seconds, as well as no windowing) to investigate the impact of this choice.

All analyses were performed in RStudio 1.4.1103 ^48^, running R 4.0.3 ^49^ and relying on the brms v2.14.4 ^50^, pROC v1.17.0.1 ^51^ and Tidyverse v1.3.0 ^52^ packages.

## Results

As shown in Figure 1a, the predicted probability of sounding happy is related to the diagnostic groups. The distribution of model predictions for the control group is stable across visits, and the distribution for patients in remission is similar to that of the control group. Predictions for the depressed group display larger variance, and are generally more sad sounding than controls.

**Figure 1:**
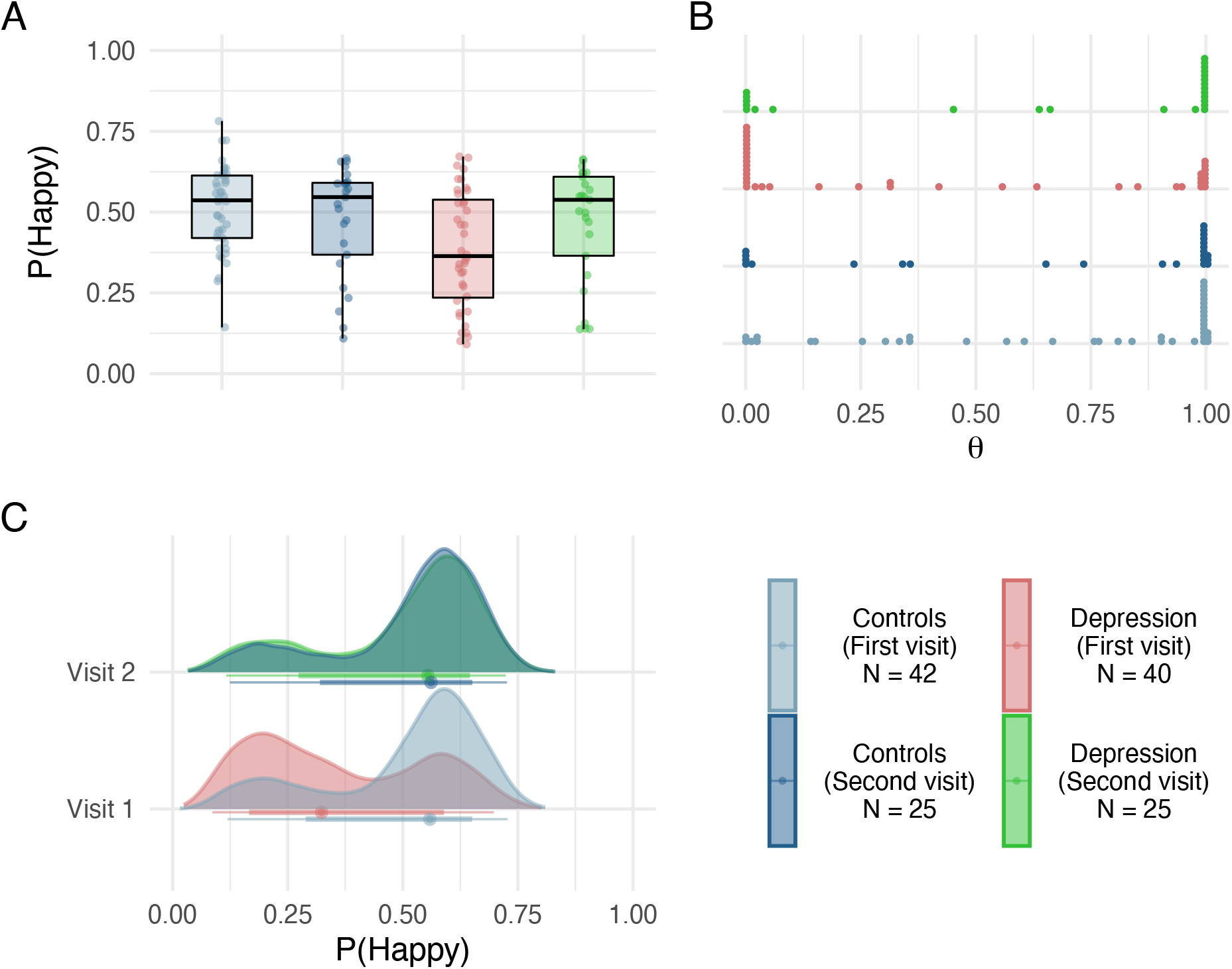
A) Distribution of mean predictions per individual at each visit. B) Probability of belonging to the ‘happy’ distribution by individual and diagnostic group. Dots show the mean mixing factor (theta) per individual. Most participants are in the extremes, which means that they are either exclusively in the happy (1) or sad distribution (0). C) Posterior probability of P(Happy) by diagnosis and visit from the multilevel Bayesian mixture model. Colored bands indicate 66% (thick line) and 95% (thin line) quantile intervals.

The multilevel Bayesian mixture model supports these observations. As seen in Figure 1b, there are two nicely separated distributions with a population level difference in the probability of belonging to the happier distribution (Figure 1c). Most participants have mixing factors (theta) very close to either 0 or 1, meaning that they are either exclusively in the *happy* or *sad* distribution. Mixing factors closer to 0.5 indicate greater uncertainty. The two Gaussian distributions identified by the mixture model are centered at 0.24 (sd=0.12) and 0.59 (sd=0.08). On the population level, depressed patients have a probability of sounding sad (theta) of 0.70 (95% CI: 0.38, 0.90), those in remission have a theta of 0.25 (95% CI: 0.07, 0.58), controls at visit 1 have a theta of 0.23 (95% CI: 0.10, 0.47), and controls at visit 2 have a theta of 0.22 (95% CI: 0.07, 0.58). The closer to 1, the larger a proportion of samples are from the *sad* distribution. The majority of depressed patients (approximately 70%) were thus identified as belonging to the *sad* distribution, whereas this number is only 22-25% for controls and patients in remission. The posterior distribution of patients in remission completely overlaps that of controls at both visits, which indicates that voice-based based symptoms of depression decrease following remission.^2^

Predictions from the SER model can discriminate between healthy controls and depressed patients obtaining an AUC of 0.71 (95% CI: 0.59-0.82) using the mean prediction for each participant. Defining the classification threshold to the one that optimizes AUC (threshold=0.38) leads to a specificity of 0.83, sensitivity of 0.55, and positive predictive value of 0.76. In other words, given an optimized decision threshold the model is able to correctly classify 83% of the control group and 55% of the depressed group, while 76% of those predicted as being depressed are correctly classified. Figure 2a shows the ROC curve for this task using both 30 second time windows and the mean prediction per participant. Predictions seem to be stable over the course of the interview when using 30 second time windows, as seen in Figure 2b^3^. The time-windowing serves to smooth small changes in the participants’ speech, and thus derives a time-independent emotional state. Further, predictions from the SER model seem sensitive to changes in the participant’s depressive state: patients in remission are more likely to be identified as controls than during their earlier depressive state (72% vs. 45% using the optimal cutoff defined by AUC).

**Figure 2:**
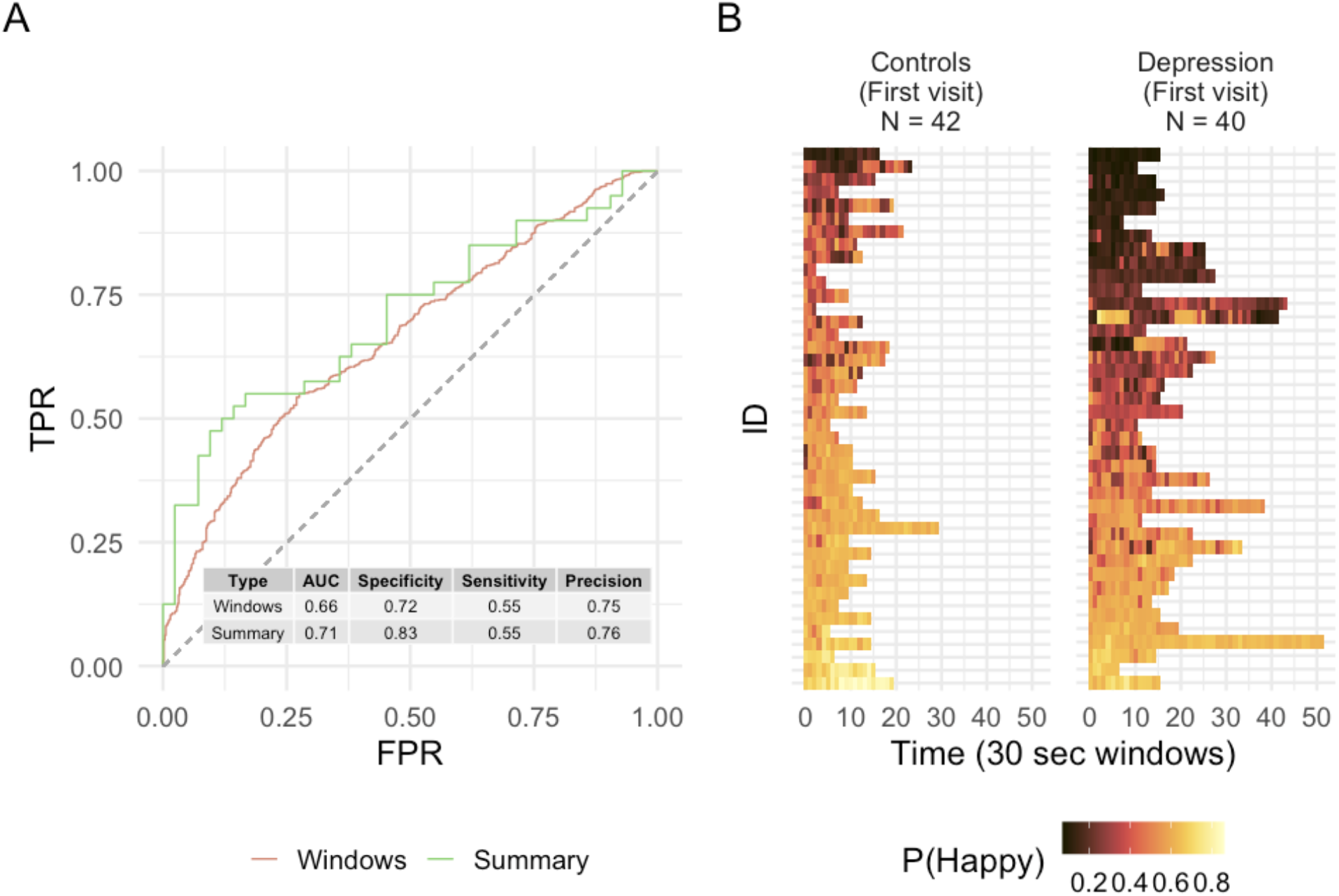
A) ROC curve displaying the separation between depression and healthy controls at visit 1. The red line is calculated on a per time-bin basis, i.e., the model is evaluated on each 30-second bin. The green line is calculated on a per participant basis, i.e. predictions are summarized using the mean for each participant. B) Heatmap showing predictions for each individual throughout the interview at visit 1 sorted in ascending order. Each row corresponds to one participant. For most participants, predictions are highly stable over the course of the interview.

The prognosis model did not find any reliable differences in voice at visit 1 between those who subsequently entered remission and those who did not. Further details are reported in Supplementary Material Figure S4.

Figure 3a shows the distribution of predictions using the different preprocessing procedures. The magnitude of the effect of background noise removal and speaker diarization differs between the groups, highlighting the importance of these procedures for consistent inferences.

**Figure 3:**
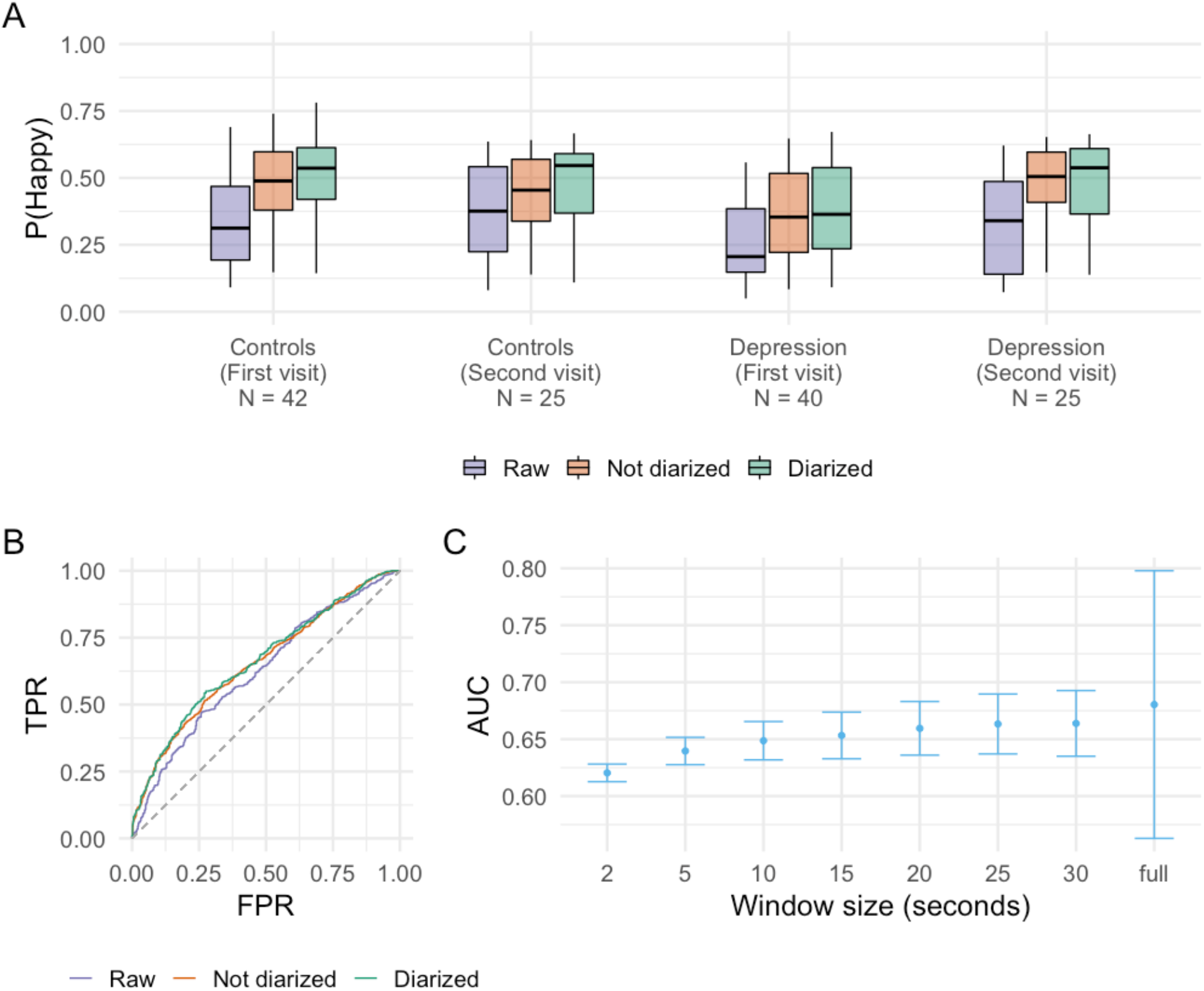
A) Difference in predictions with different preprocessing strategies. Both the diarized and non-diarized datasets were cleaned from background noise. B) ROC-curve for the task of predicting depression or control at visit 1 with different preprocessing strategies. Using 30 second windows. C) AUC for the task of predicting depression or control at visit 1 using different timewindow sizes at a per window basis. Error bars display 95% bootstrapped confidence intervals. Note that the last bar presents higher uncertainty as it is evaluated on a per-participant basis where the others are evaluated on a per-time-window basis.

The effect of the preprocessing steps on the model’s ability to discriminate between healthy controls and depressed patients is visualized in Figure 3b. The lowest AUC is obtained using raw data (AUC: 0.63, 95% CI: 0.60-0.66), followed by background noise removal and no speaker diarization (AUC: 0.66, 95% CI: 0.63-0.68), with data using both speaker diarization and background noise removal obtaining the highest AUC (AUC: 0.66, 95% CI: 0.63-0.69).^4^

Correspondingly, the model following the BEST approach ^47^ found marked differences between models trained on data with different levels of preprocessing, although with large uncertainty estimates. The model trained on raw data had the lowest effect size for the difference between controls and depression (Cohen’s d 0.39, 95% CI: 0.08-0.71), background noise removal but no speaker diarization was in the middle (Cohen’s d 0.49, 95% CI: 0.18-0.77), and background noise removal with speaker diarization had the largest effect size (Cohen’s d 0.55, 95% CI: 0.24-0.86). Although the effect of speaker diarization is not as extreme as background noise, performing speaker diarization improves model performance.^5^

As shown in Figure 3c, AUC gradually increases with larger time-window sizes, however there seems to be a ceiling effect around windows of 20 seconds. The same figure shows that using the whole recording (no windowing) provides superior performance than assessing on a per-window basis. However, taking the mean prediction per participant using 20-30 second windows was found to be better than using the whole recording (no windowing: AUC: 0.68, 95% CI: 0.56, 0.80; mean of 30 second windows: AUC: 0.71, 95% CI: 0.59, 0.82).

## Discussion

This study set out to investigate three main aims: 1) whether a speech emotion recognition model is useful for predicting depression 2) whether these predictions can be used for assessing changes in voice following remission, and 3) how much preprocessing steps impact the quality of model predictions.

We found that the speech emotion recognition model was able to accurately discriminate between healthy controls and depressed patients, obtaining an AUC of 0.71 (95% CI: 0.59-0.82). While voice patterns during the course of the disorder did not predict whether a patient would remit in the future, remission itself could be observed in the voice. Indeed, a remission effect was observed, as the voice of patients in remission was indistinguishable from that of healthy controls and was credibly more *happy* sounding than during the disease. Model predictions were stable over the course of the interviews when using a 30-second time window, indicating that robust, long-term representations of voice are captured. Pre-processing steps had an impact on the model predictions: background noise had a large effect on model performance, and must be controlled for making meaningful inferences. Removing speech from the interviewer (speaker diarization) did not reliably affect model performance, however, it slightly increased effect size. Model performance was found to increase with larger window sizes at the expense of increased variance, and to stabilize around 20-30 seconds.

### Heterogeneity in Depression

Although we found a credible difference in the voice of depressed patients and healthy controls there seem to be two subpopulations of people with depression. Approximately 30% of the patients in the depressed group were predicted as having similar emotional valence (from sad to happy) of their voices as the control group, whereas the remaining 70% have markedly more ‘sad’ sounding voices (see Figure 2). This large variability potentially arises from the fact that MDD is a highly heterogeneous disorder ^53,54^, and patients might therefore express disparate symptoms while still falling under the umbrella of MDD ^45^. To partly account for this, MDD can be subdivided into a melancholic, anxious, and atypical type based on distinct symptomatic profiles ^55^. The melancholic subtype might be of particular relevance for depression detection from speech, as it is characterized by severe anhedonia, psychomotor disturbance, and neurovegetative symptoms ^56^. Investigating whether specific depression subtypes are better captured by speech analysis than others is a field of further research. However, approximately 44% of patients can not be assigned a specific subtype, and only 15% have the melancholic subtype ^55^. As a consequence, significant unexplained heterogeneity between patients remains and a more granular perspective based on specific symptoms might be better poised at describing this heterogeneity ^57^. Models of depression from speech are inherently constrained by this factor, which underscores that the primary area of application for such systems should be screening and disease monitoring, not diagnosis.

### Generalizability

The main focus of our work was to improve generalizability and robustness of depression detection from voice. In this regard, factors relating to intrapersonal variability, method of speech elicitation, and language must be discussed.

The intrapersonal variability among participants was heterogeneous. Within each interview, the probability of sounding happy varied on average from 5.7 to 7.9 percentual points depending on the diagnostic group. The distribution of this value was long-tailed, meaning that predictions were relatively stable for the majority of participants, with a minority being very variable. Though clear trends were visible on the group aggregate level, the extent to which each participant’s voice changed between visits differed markedly. Parts of the variability might stem from participants being in a different emotional state, from slightly different recording settings, or from different changes in depressive symptom profiles across visits. For example, one depressed patient might have started with a low number of symptoms influencing the voice and therefore not show a large change at visit two, whereas another might be in the opposite situation which would lead to a large change in predictions at the second visit. To increase robustness of the method, we advise practical applications to perform multiple recordings over multiple days. Further, to better understand the symptom profiles and patient cohorts which might benefit from voice-based depression detection systems, further studies should strive to include variables relating to symptom expression.

Several previous studies have found the method of speech elicitation to impact the patterns extracted from speech ^58,59^. Patterns of pathological voice are expressed to a greater extent in more social and cognitively demanding tasks such as free speech than in read speech or vocal exercises ^9,16^. A clinical interview can be considered an extremely social and cognitively demanding task, and might therefore provide an exceptionally strong signal for detecting depression from voice. Whether our model works equally well on voice elicited from other tasks remains to be tested. An advantage of our data collection as compared to gathering data more naturalistically from e.g. phone samples or in participants’ home, is that the context is highly controlled. This means that biases related to background noise are greatly reduced. For instance, there could be systematic differences in the noise level, or number of people in the background between the two populations which might confound a model. However, it would be advisable to extend this study to such as dataset in the future to further assess generalizability.

Our model was trained to predict emotion in voice from English and German speech and validated on Danish speech for the task of detecting depression, thereby generalizing to new participants, tasks, and even to a new language. As a consequence, our results are highly likely to generalize to other Germanic languages recorded in similar ways and similar samples. This is valuable for clinical implementations, as models need to handle a variety of languages, dialects, and accents. Further, there is no risk of overfitting to the data, as our model’s hyperparameters were optimized for SER, not prediction of depression. However, our model was only trained and tested on Germanic languages which leaves the extent of generalizability across language families unknown. The finding that emotional valence of speech (from sad to happy) from patients in remission is similar to healthy controls increases the credibility of our model, and suggests that acoustic features of speech could be used as an effective marker for depression. However, it should be noted that we only had access to 25 patients in remission and that we were not able to make comparisons with the group who remained depressed.

### Effect of Preprocessing

This paper sought to investigate the effect of different preprocessing methods for producing reliable predictions, namely background noise removal, speaker diarization, and window size.

Background noise was found to drastically impact model performance by differentially distorting predictions based on the level of noise in the recordings. This has major implications for clinical implementations as recordings must be made under relatively noise-free conditions to avoid excessive false positives. For screening and monitoring, participants can be advised to perform the audio recordings in quiet surroundings, however automated quality checks and noise reduction methods are likely necessary.

Speaker diarization had some effect on model performance, but not to the same extent as noise removal. The audio recordings used in this study consisted of interviews conducted by the same psychologist with all the participants. In such a dyadic setting, the psychologist is likely to align their way of speaking with that of the participants thereby decreasing the effect of speaker diarization ^60-62^. However, audio recorded in environments with multiple speakers or in more naturalistic settings (e.g. with sounds from people in adjacent rooms) will likely present more severe confounds without a speaker diarization process. The extent to which speaker diarization will impact predictions will naturally vary with how large a proportion of speech the interviewer produces. The type of interview used in this study (Indiana Psychiatric Illness Interview) tends to have a relatively low proportion of interviewer speech compared to other psychiatric interviews. This means that other interview types might see greater benefit from speaker diarization.

Larger window sizes, i.e the size of the time bin used for summarization of acoustic features, afforded better predictions until a ceiling effect at window sizes of 20-30 seconds of speech. Using the whole recording without windowing performed marginally better than windowing when evaluating on a per-window basis. However, when summarizing the prediction for each window into a single prediction per participant, the windowed summarization outperformed the whole recording, again with a ceiling effect at window sizes of 20-30 seconds. This suggests that recordings as short as 20-30 seconds of speech might be enough to provide high quality predictions, but longer recordings, if available, will slightly improve performance.

### Limitations

The results of our study need to be interpreted in light of the following limitations. First, this study mainly served to investigate the usefulness of directly applying transfer learning from SER to the task of predicting depression under optimal conditions. In this regard, manual preprocessing procedures such as background noise removal and speaker diarization and corresponding quality checks were taken, but are not feasible for clinical implementations to the same extent. A large part of the preprocessing pipeline can be automated, but performance is likely to suffer without a human in the loop.

Second, given that the focus of our study has been more on generalizability than predictive performance, we decided to only train our model on MFCCs although several other acoustic features have been found predictive of depression ^9^. As reviewed, MFCCs have several desirable properties for generalization, whereas features such as pitch are highly gender dependent and therefore might harm generalizability. However, once larger and higher quality datasets of depressed speech become available, it might be beneficial to include features more specific to depression. In a similar vein, the model proposed here could easily be extended by training depression-specific experts and adding to the ensemble. Further, owing to the success of transformer-based neural network models in fields such as Natural Language Processing, it might prove beneficial to use speech models such as wav2vec 2.0 ^63^, or its multilingual variant ^64^, for SER and depression detection.

Third, our models are sensitive to the preprocessing steps and recording environment. Data must be carefully cleaned to ensure consistent noise profiles and inferences. Though our model was robust against speech from the interviewer, background noise from other people might further influence model predictions.

Fourth, though our model was validated on a relatively larger dataset than many other studies in this field, there is a pronounced need for larger, high-quality, longitudinal datasets from diverse languages. Current publicly available databases often contain a low number of participants, sparse information on symptoms and demographic characteristics, and are primarily in English or other Germanic languages. Longitudinal studies following the same patient group over the course of their treatment could provide insights into the effectiveness and robustness of voice-based depression measures and potentially cast light on which subpopulation of patients the models work best for. Datasets from more diverse languages are required to assess cross-lingual performance. Our study provides an attempt at this, by training and testing on different languages.

Fifth, it is unknown how our model would predict speech produced by patients with other disorders known to cause abnormal speech such as bipolar disorder, schizophrenia, Parkinson’s, or autism spectrum disorder, or from other common disorders such as anxiety. In this study, we focus on a proof of concept: can we use models trained on non-clinical data to assess clinical data, and therefore limit ourselves to the case of MDD. Given the lack of knowledge of the model’s specificity to MDD, the main clinical application of the model in its current form would likely be tracking of remission status or effect of treatment in patients already diagnosed with MDD. Cross-diagnostic and symptom-based approaches are important extensions for future work.

Last, from a clinical perspective it might not be desirable to have a model that classifies controls and remitted patients similarly, as remitted patients are more likely to relapse to MDD ^65^. Future work is encouraged to develop more sensitive models, for instance by finetuning on clinical data or specifically target risk of relapse or remission.

### Conclusions

Voice-based systems have the advantage of being less prone to biases related to self-reports and human ratings, and can be used remotely, cheaply, and non-invasively. Successful implementation of voice-based depression screening and monitoring has potential for providing earlier diagnosis and a more granular view of treatment effect, thereby facilitating improved prognosis of major depressive disorder. To reach this aim we showed the potential of transfer learning to identify the presence of depression, and identified conditions of applicability: at least 30 seconds of voiced speech, strong attention to background noise and recording conditions, but no huge impact of diarization.

## Appendix

### Speech Emotion Recognition with Mixture of Experts

We made use of the model trained in Sechidis et al. ^1^ for SER and briefly describe it here for completeness. The SER model was trained on the CREMA-D ^2^, RAVDESS ^3^, and EMO-DB ^4^ datasets and validated on EMOVO ^5^, TESS ^6^, and SAVEE ^7^. Each corpus contains recordings from professional actors who repeat sentences while changing which emotion they convey. MFCC coefficients were extracted from each recording (10 ms windows), and summarized using the 11 descriptive statistics mentioned in the Methods section (mean, variance, kurtosis, skewness, mode, IQR, percentiles 10th, 25th, 50th, 75th, and 90th) per utterance.

For each training dataset, a gradient boosted decision tree model was fitted using Cat-Boost. The optimal hyperparameters for each model were estimated using subject-wise crossvalidation stratified by gender. The three models were combined in a Mixture of Experts (MoE) architecture using the Mahalanobis distance as similarity metric. In practice, this entails that for each new sample to predict, the prediction from each constituent model is weighted in terms of how similar the new datapoint is to the model’s training data.

To assess the effectiveness of the MoE, its performance was tested against each constituent model on their own, as well as a model trained on pooled data from all three datasets. Performance was assessed on the EMOVO, TESS, and SAVEE dataset on which the MoE was found to achieve superior performance (average accuracy: 0.84)^1^.

### Model Building

The Bayesian mixture model trained to assess the difference in the probability of sounding happy from the interaction between diagnosis and visit was trained using the *brms* R package ^8^. The probability of sounding happy was transformed to log odds to improve model fit, and subsequently modelled as a mixture of two Gaussian distributions. Weakly regularizing priors were used for all parameters:

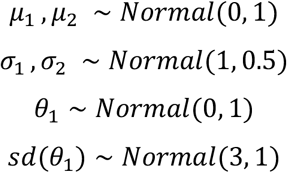

The model was trained for 4000 iterations, including 1000 warmup iterations, on 4 chains, with adapt_delta set to 0.99 to improve convergence. All 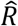 ^9^ were below 1.001 and chains were visually inspected for convergence with no obvious issue being found. See Supplementary Material Figure S6 and S7 for prior and posterior predictive checks and posterior update plots.

The Bayesian multilevel model trained to assess the difference in the probability between the diagnostic groups varying by level of preprocessing were also trained using the *brms* R package. One model was fit for each dataset (with different levels of preprocessing), with the same settings with regards to priors and samples. The models were fit following Kruschke ^10^, and modelled as a T-distribution. The priors for the betas were set as Gaussians with mean 0.5, and standard deviation 0.5, bounded at 0 and 1. Prior for the *nu* parameter for the normality of the T-distribution was an exponential with parameter 1/29 following Kruschke’s ^10^ recommendation. Models were run for 6000 iterations on 4 chains, and passed all checks for convergence.

## Footnotes

1 See Appendix for further details on model building and assessment of the quality of the model fit.

2 See Supplementary Figure S1 for changes in P(Happy) between visits for each participant.

3 See Supplementary Figure S5 and Supplementary Table S4 for individual and group level standard deviations.

4 See Supplementary Table S2 for more performance metrics.

5 See Supplementary Table S3 and Supplementary Figure S3 for full model report.

