## Supplementary materials for "A Generalizable Speech Emotion Recognition Model Reveals Depression and Remission"

### Supplementary Material

*Supplementary Table S1: Duration of speech after noise removal, and interview length.*

| | | Participant speech duration | | Total speech duration | | Interview length | |
|---|---|---|---|---|---|---|---|
| Diagnosis | Visit | Mean | SD | Mean | SD | Mean | SD |
| Controls | 1 | 6.39 | 2.82 | 10.19 | 3.64 | 16.23 | 7.18 |
| Controls | 2 | 5.94 | 2.40 | 8.46 | 2.38 | 13.29 | 3.12 |
| Depression | 1 | 10.20 | 5.03 | 14.75 | 5.40 | 28.23 | 9.43 |
| Depression | 2 | 8.62 | 6.07 | 12.20 | 6.68 | 21.43 | 12.29 |

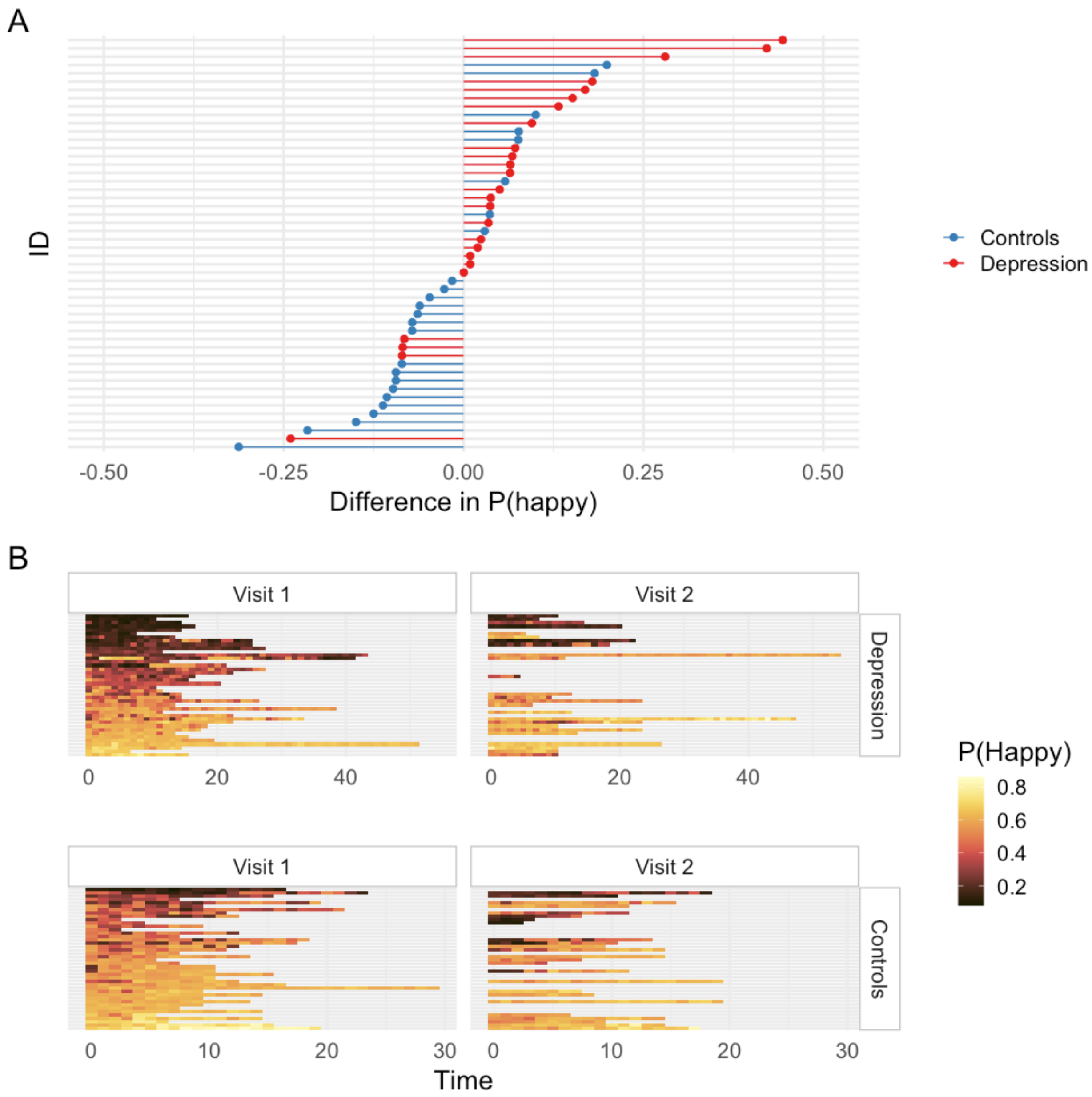


*Supplementary Figure S1: A) Lollipop chart displaying the mean change in predictions from visit 1 to visit 2. Each row corresponds to one participant. In general, patients in remission sound happier at visit 2 than at visit 1. B) Heatmap showing predictions throughout the interview for all participants at both visits.*

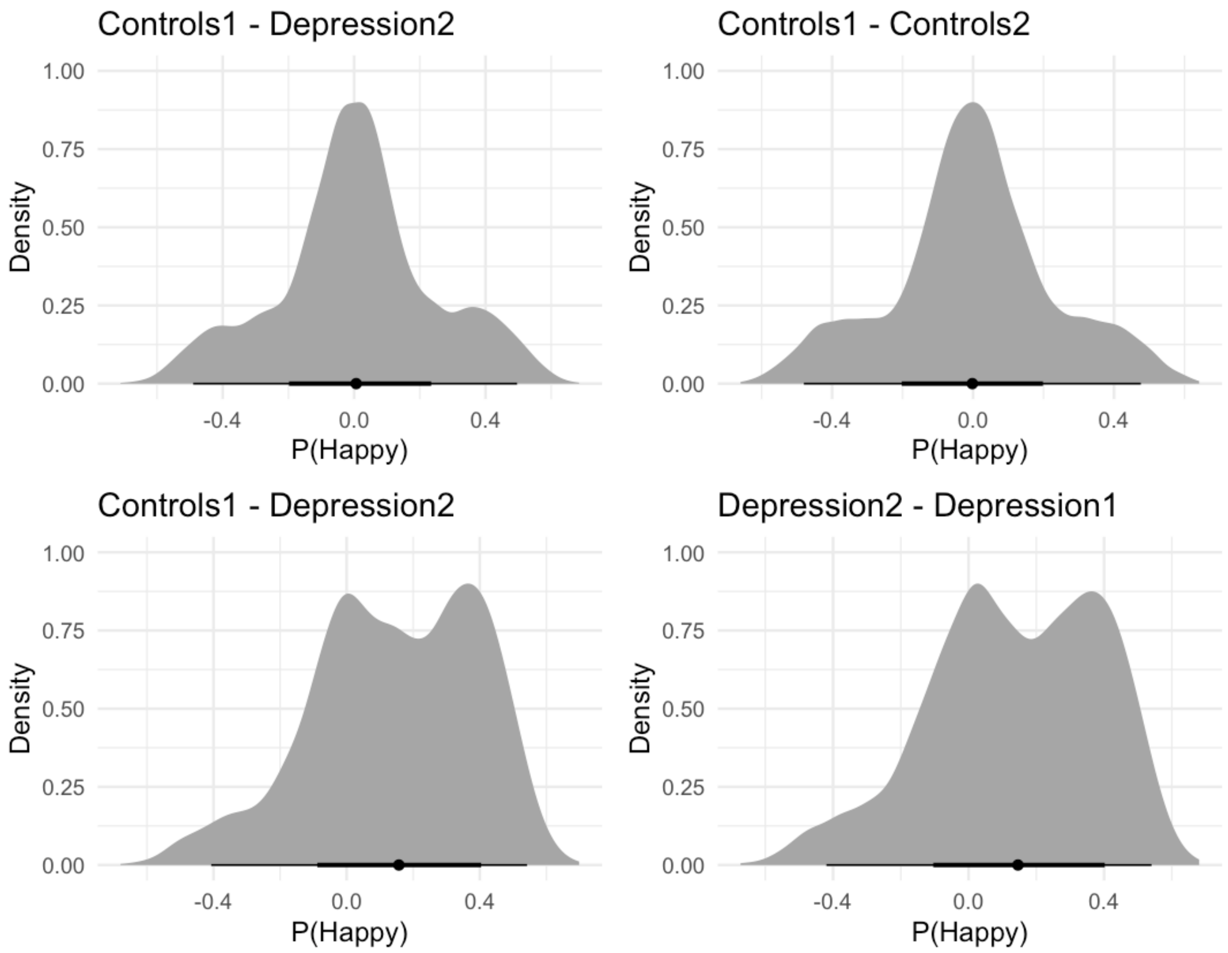


*Supplementary Figure S2: Difference in estimated P(Happy) between the diagnostic groups using the mixture model.*

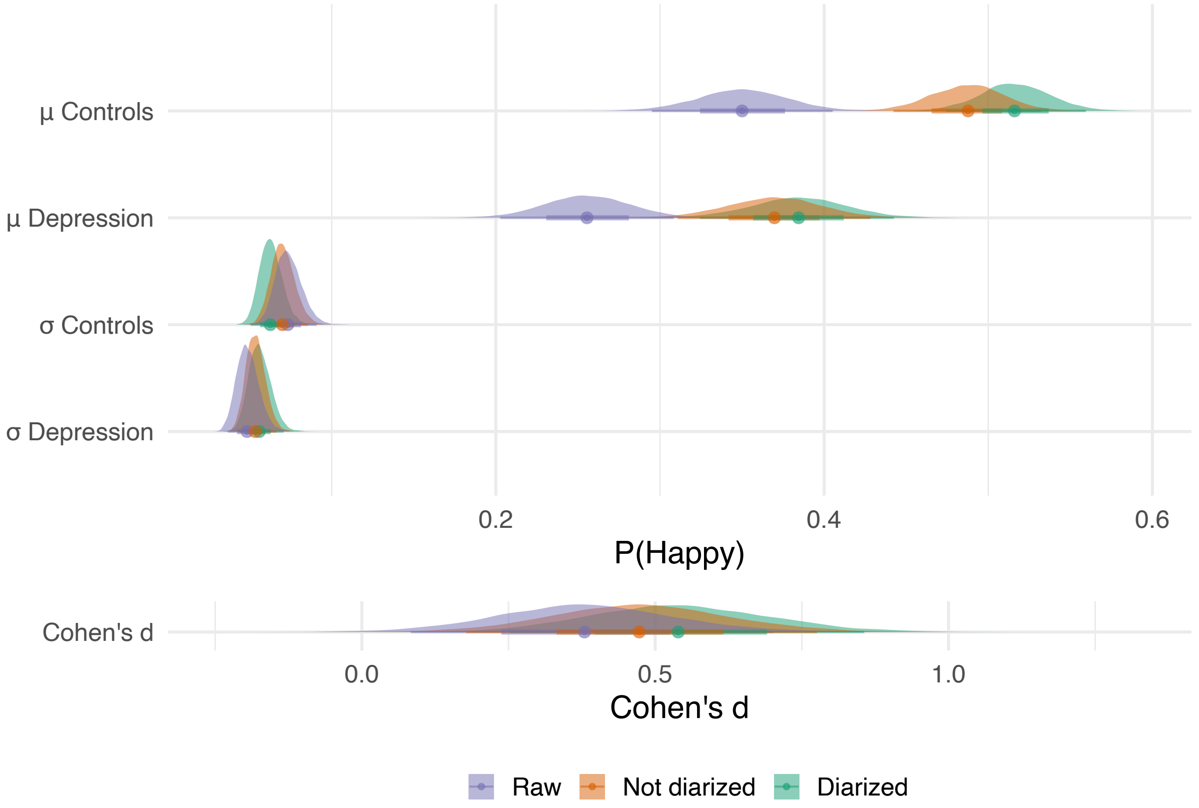


*Supplementary Figure S3: Results from the multilevel Bayesian model (Kruschke, 2012) assessing the difference between healthy controls and depression at visit 1 using different preprocessing methods. Both the diarized and non-diarized datasets were cleaned from background noise. Note the difference in x-axis for Cohen's d.*

*Supplementary Table S2: AUC on the different datasets using 30 second windows.*

| Data | AUC | AUC CI | Threshold | Specificity | Sensitivity | Precision |
|---|---|---|---|---|---|---|
| **Summary** | | | | | | |
| Diarized | 0.71 | 0.59-0.82 | 0.38 | 0.83 | 0.55 | 0.76 |
| Not diarized | 0.69 | 0.57-0.8 | 0.36 | 0.81 | 0.55 | 0.73 |
| Raw | 0.67 | 0.55-0.79 | 0.16 | 0.95 | 0.36 | 0.88 |
| **Windows** | | | | | | |
| Diarized | 0.66 | 0.63-0.69 | 0.44 | 0.72 | 0.55 | 0.75 |
| Not diarized | 0.66 | 0.63-0.68 | 0.37 | 0.73 | 0.51 | 0.72 |
| Raw | 0.63 | 0.6-0.66 | 0.19 | 0.74 | 0.47 | 0.74 |

*Supplementary Table S3: Full tabular results from the multilevel Bayesian model (Kruschke, 2012) assessing the difference between healthy controls and depression at visit 1 using different preprocessing methods.*

| Dataset | Estimate | Mean 95% QI |
|---|---|---|
| **Cohens's d** | | |
| Raw | 0.39 | 0.08-0.71 |
| Not diarized | 0.48 | 0.18-0.77 |
| Diarized | 0.55 | 0.24-0.86 |
| **Diff. means** | | |
| Raw | 0.10 | 0.02-0.17 |
| Not diarized | 0.12 | 0.04-0.19 |
| Diarized | 0.13 | 0.06-0.21 |
| **Diff. sigma** | | |
| Raw | 0.03 | 0.01-0.05 |
| Not diarized | 0.02 | 0-0.03 |
| Diarized | 0.01 | -0.01-0.02 |
| **µ Controls** | | |
| Raw | 0.35 | 0.3-0.41 |

| | | |
|---|---|---|
| Not diarized | 0.49 | 0.45-0.53 |
| Diarized | 0.52 | 0.47-0.56 |
| **μ Depression** | | |
| Raw | 0.26 | 0.2-0.31 |
| Not diarized | 0.37 | 0.31-0.43 |
| Diarized | 0.38 | 0.32-0.44 |
| **σ Controls** | | |
| Raw | 0.07 | 0.06-0.09 |
| Not diarized | 0.07 | 0.06-0.09 |
| Diarized | 0.06 | 0.05-0.08 |
| **σ Depression** | | |
| Raw | 0.05 | 0.04-0.06 |
| Not diarized | 0.05 | 0.04-0.07 |
| Diarized | 0.06 | 0.04-0.07 |

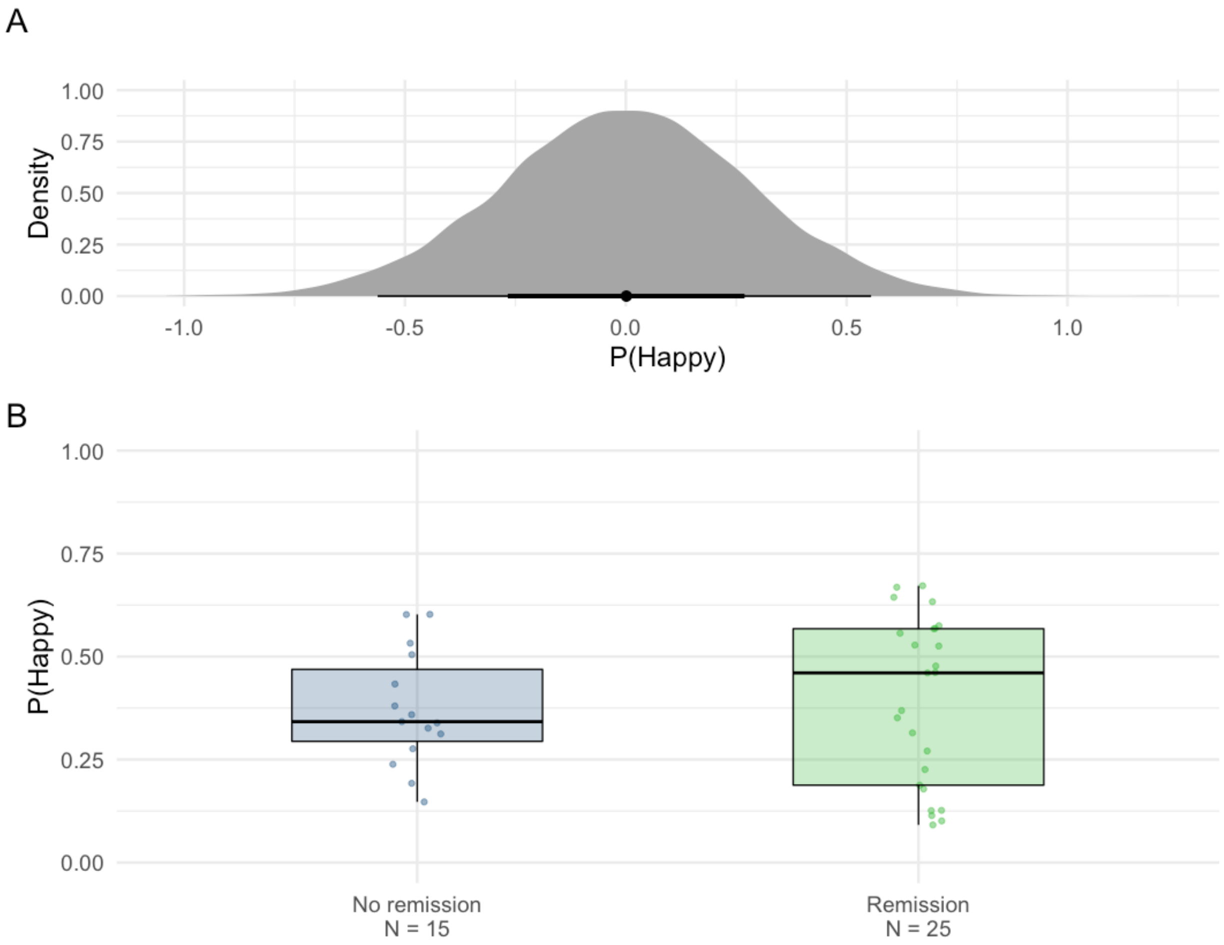


*Supplementary Figure S4: Predicting who will enter remission (prognosis). A) estimated difference in the probability of sounding happy at visit 1 for those who will enter remission and those who won't, B) distribution of predictions at visit 1 for those who will enter remission and those who won't.*

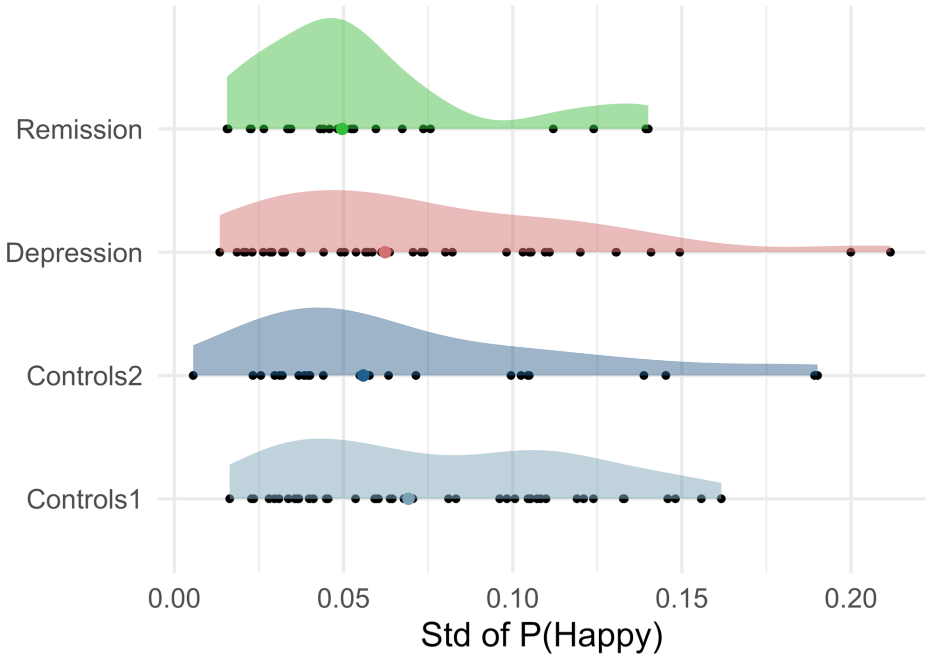


*Supplementary Figure S5: Stability of predictions with 30 seconds windows. Standard deviation of P(Happy) per participant. Colored dots mark the mean.*

*Supplementary Table S4: Mean standard deviation of P(Happy) and corresponding standard deviation per diagnostic group.*

| Diagnosis | Mean std P(Happy) | Std of mean std P(Happy) |
|---|---|---|
| Controls1 | 0.079 | 0.042 |
| Controls2 | 0.071 | 0.051 |
| Depression | 0.074 | 0.048 |
| Remission | 0.057 | 0.036 |

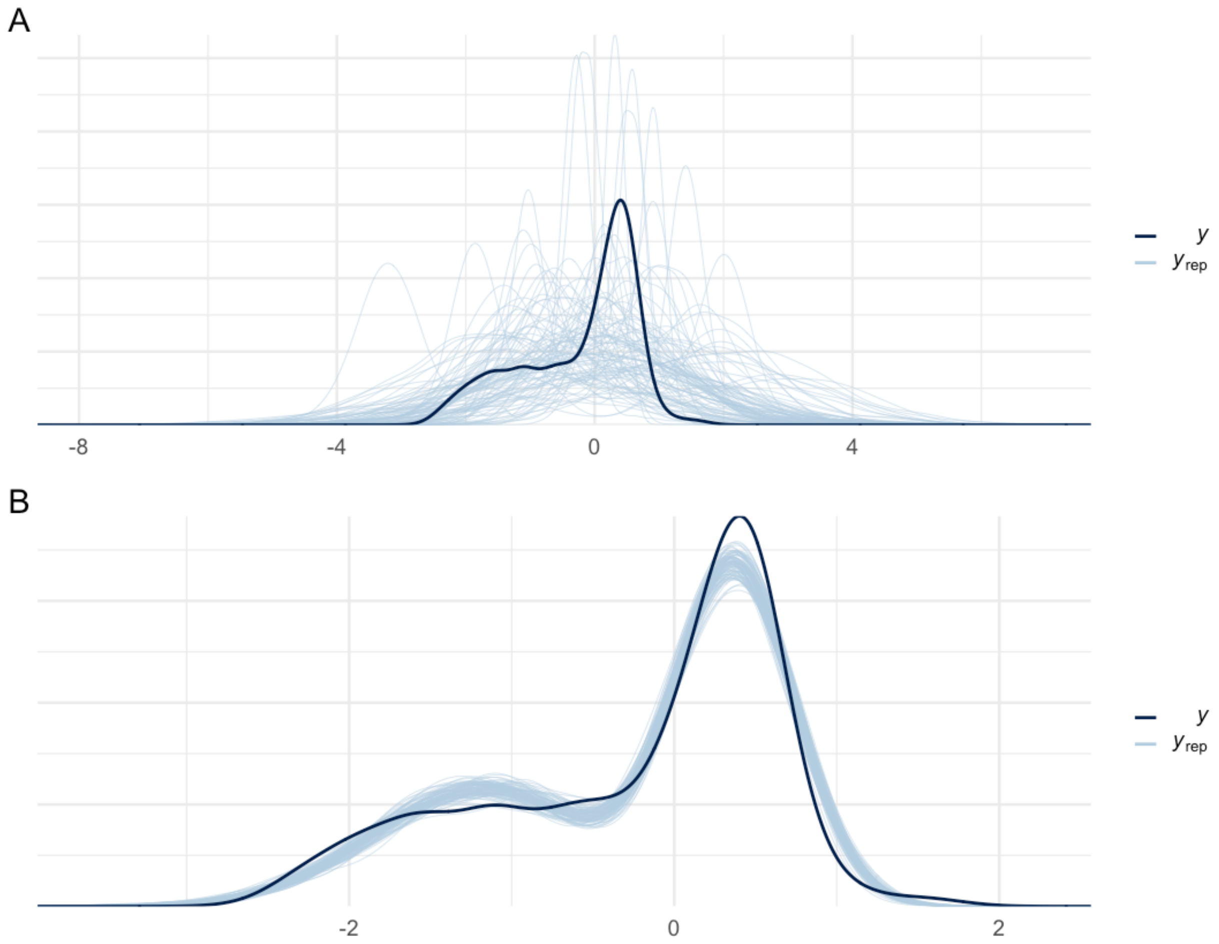


*Supplementary Figure S6: Prior and posterior predictive checks for the mixture model. A) Prior predictive check, B) posterior predictive check.*

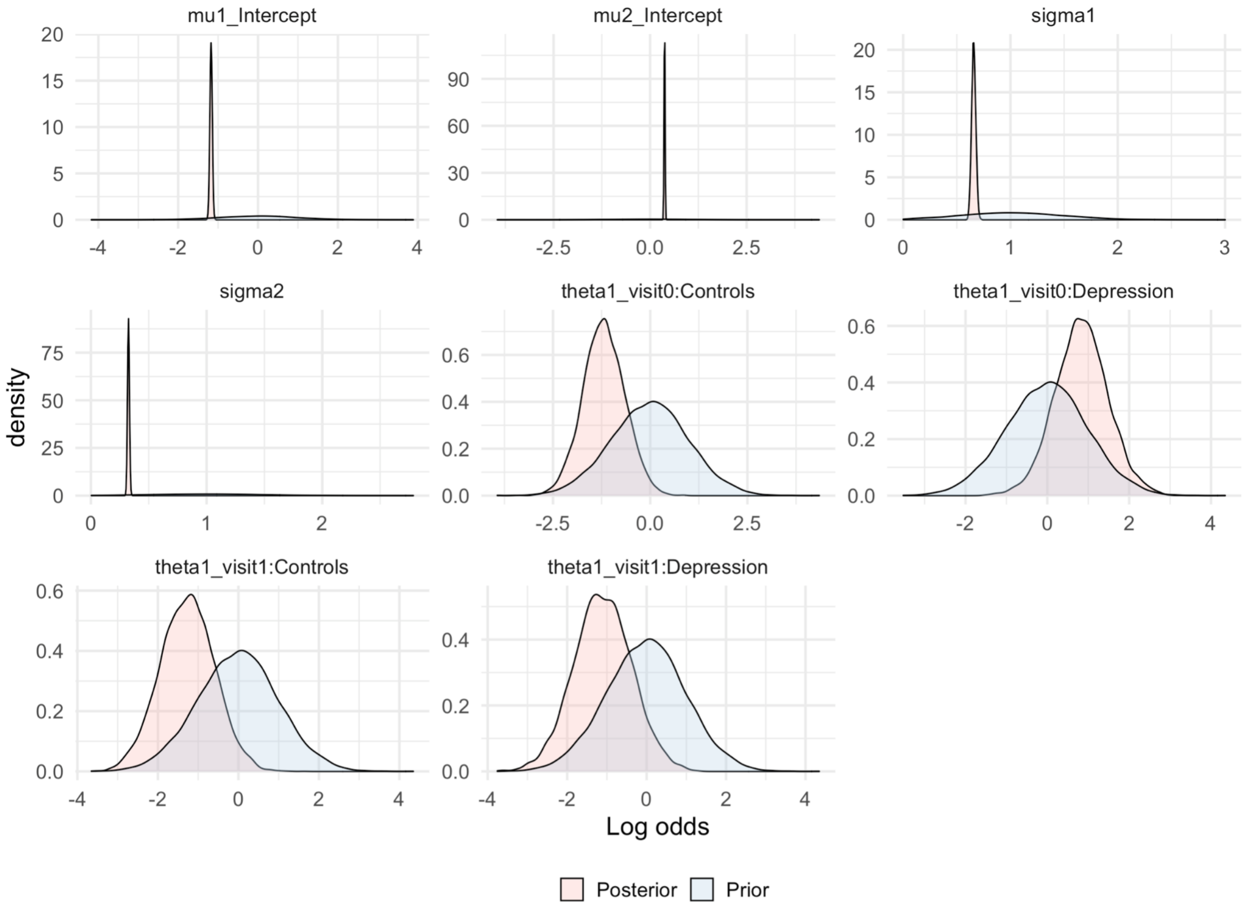


*Supplementary Figure S7: Posterior updates for the mixture model. Note that the values are on a log odds scale.*

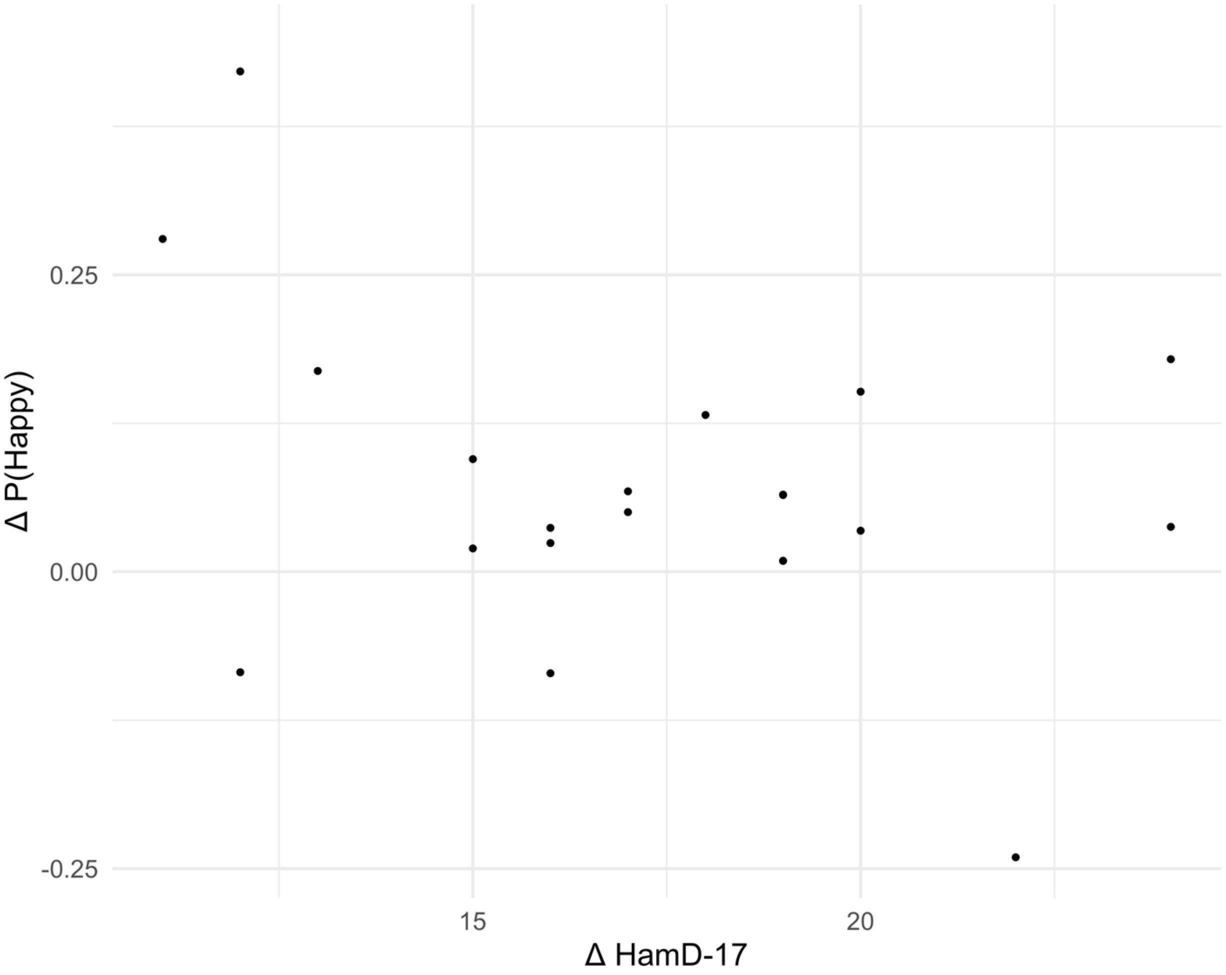


*Supplementary Figure S8: Difference in P(Happy) from first to second visit as a function of the difference in the 17 item Hamilton Rating Scale for Depression from the first to the second visit for depressed patients only. The larger the value on the x-axis, the greater the drop in Hamilton score, and the larger the value on the y-axis the larger the P(Happy) at second visit.*
